# Your turn, my turn. Neural synchrony in mother-infant proto-conversation

**DOI:** 10.1101/2022.09.09.507239

**Authors:** Trinh Nguyen, Lucie Zimmer, Stefanie Hoehl

## Abstract

Even before infants utter their first words, they engage in highly coordinated vocal exchanges with their caregivers. During these so-called proto-conversations, caregiver-infant dyads use a presumably universal communication structure - turn-taking, which has been linked to favourable developmental outcomes. However, little is known about potential mechanisms involved in early turn-taking. Previous research pointed to interpersonal synchronisation of brain activity between adults and preschool-aged children during turn-taking. Here, we assessed caregivers and infants at 4-6 months of age (N=55) during a face-to-face interaction. We used functional-near infrared spectroscopy hyperscanning to measure dyads’ brain activity and microcoded their turn-taking. We also measured infants’ inter-hemispheric connectivity as an index for brain maturity and later vocabulary size and attachment security as developmental outcomes potentially linked to turn-taking. The results showed that more frequent turn-taking was related to interpersonal neural synchrony, but the strength of the relation decreased over the course of the proto-conversation. Importantly, turn-taking was positively associated with infant brain maturity and later vocabulary size, but not with later attachment security. Taken together, these findings shed light on mechanisms facilitating preverbal turn-taking and stress the importance of emerging turn-taking for child brain and language development.

## Introduction

Long before acquiring verbal language, human infants engage with their caregivers in reciprocal vocal exchanges, so-called proto-conversations [1–6], which have a strikingly similar temporal structure to later verbal conversations [7]. Both types of exchanges are characterised by a rapid back-and-forth (i.e., turn-taking) between the interaction partners, with short lags between the turns and limited overlaps. This temporal structure is documented across many languages and cultures, suggesting that the turn-taking system might be a universal feature of human communication [8]. In the current study, we set out to explore the early origins of turn-taking, rooted in the infant-caregiver interaction. What are the mechanisms of early turn-taking, and what are its functions in preverbal human infants?

In human adults, turn-taking enables highly efficient verbal conversations [9]. Across languages, response latencies between speakers are remarkably short (around 200ms on average), implying that speakers must predict the speech act of their interlocutor while still listening, and they must plan their response before their partner’s speech offset [10]. Combined with rare overlaps of partners speaking simultaneously, these cognitive processes ensure a fluid conversation and highly efficient exchange of information. Turn-taking is a fundamentally cooperative endeavour as partners must pay close attention to each other to enable such fine-grained mutual coordination. Accordingly, shorter response latencies have been associated with greater feelings of connection between interaction partners [11].

In early human ontogeny, infants as young as eight weeks of age actively engage in turn-taking in vocal interactions with their caregivers [1]. These early exchanges already display a remarkable level of temporal coordination [12]. In the first six months after birth, infants’ response latencies are quite similar to latencies observed in adult conversations [2]. The early ontogenetic onset of turn-taking in human social exchanges and the structural similarity of preverbal proto-conversations with later verbal conversations beg the question of whether similar mechanisms are involved in early and later turn-taking. The objective of the current study is twofold: (i) We aim for a better understanding of the mechanisms of early turn-taking exchanges between young infants and their mothers, and (ii) we want to assess potential links between early dyadic turn-taking and later developmental outcomes, in particular infant attachment quality and vocabulary size.

### Mechanisms of early turn-taking

As turn-taking depends on precise temporal coordination, one key candidate underlying mechanism is the mutual entrainment and synchronisation of endogenous neural oscillators in the brains of the conversation partners [13]. According to this view, a listener’s neural oscillations become entrained to the rhythms of their partner’s speech and vice versa, leading to mutual synchronisation of brain rhythms [14] which enables fine-tuned conversational turn-taking. Indeed, ample empirical evidence shows that listeners’ neural oscillations entrain to perceived speech rhythms in adults [15,16] and infants [17,18]. Using hyperscanning, meaning simultaneous measurements of brain activity of at least two participants, interpersonal synchronisation of brain rhythms has been documented in verbal conversations between adults [19,20] and between caregivers and their preschool-aged children [21]. The growing body of research underscores the involvement of temporal and especially frontal brain regions in neural synchrony during a conversation [22]. Synchrony in these brain regions has been related to precise mutual predictions, which are integral cognitive processes to turn-taking. Indeed, neural synchronisation during mother-child conversation increased more steeply when dyads took more turns in their conversation [21].

Importantly, we know little about when the relation between interpersonal neural synchrony and turn-taking emerges during development. Gaining insights into the functional role of interpersonal neural synchrony is important as it could pave the way for interventions promoting neural synchrony and its related benefits [23,24]. Furthermore, if turn-taking is linked to neural synchrony already in infant-caregiver interactions this might point to neural synchrony as a potential shared mechanism underlying turn-taking across ontogeny. Given that brain rhythms undergo protracted development in humans [25] mutual adaptation of rhythmic brain activities, resulting in neural synchrony during turn-taking, could be an important factor in shaping the development of infant brain dynamics [26]. In the present study, we, therefore, tested whether interpersonal neural synchrony relates to turn-taking already in mother-infant dyads. We hypothesised that turn-taking is linked to higher mother-infant neural synchrony, which should increase over the course of a free proto-conversation. Specifically, we predicted that higher interpersonal neural synchrony relates to more frequent and faster turns, indicating higher turn-taking quality [7] (Hypothesis 1).

### Links between early turn-taking and developmental outcomes

As infants continue to partake in early proto-conversations, including turn-taking, they become more competent at turn-taking and increasingly initiate turn-taking sequences [1]. We suggest that individual differences in becoming more mature participants in proto-conversations might be reflected in variability in infants’ functional brain network connectivity [27]. Especially functional connectivity in the homologous inter-hemispheric network, which is considered a marker for neural maturity [27,28], could be linked to infants’ early turn-taking abilities. According to previous literature, these abilities should include more frequent turn-taking initiated by the infant as well as faster infant responses to turns initiated by the mother [1,2,7]. In the present study, we measured infant maturity at a neural level using inter-hemispheric connectivity and linked it to turn-taking behaviour between mothers and their infants. We hypothesised that more frequent infant-initiated turn-taking and, potentially, turns with shorter response latencies are associated with higher inter-hemispheric connectivity (Hypothesis 2).

Early turn-taking has been associated with developmental benefits in language processing and acquisition [10,29]. For instance, higher rates of turn-taking have been related to advanced language skills [30] and larger vocabulary in 2-year-olds [31]. In contrast, weaker vocal coordination of preverbal infants has been linked to poorer cognitive outcomes [12]. In line with these findings, we hypothesised that more frequent turns between mothers and their infants and turns with shorter response latencies are linked to larger vocabulary at two years of age (Hypothesis 3).

Next to infants’ individual development, turn-taking in mother-infant proto-conversations is assumed to be one fundamental feature of interpersonal connectedness [11,22] and the basis for parent-child bonding [32]. More specifically, insecurely-attached children, a measure of parent-child relationship quality [33], show longer response latencies during preverbal turn-taking than securely attached children. Still, there is little evidence on the role of preverbal turn-taking for attachment development, even though there is ample evidence for the role of behaviourally coordinated interactions for attachment security. In the present study, we thus examined whether turn-taking patterns in mother-infant proto-conversation can inform us about infants’ attachment development. Accordingly, we assumed that more frequent turns and faster turns, i.e., shorter response latencies, are related to higher attachment security (Hypothesis 4).

## Methods

### Participants

Fifty-five mother-infant dyads (25 female infants; M age = 145.36 days, SD = 16.93 days; range = 121-180 days) were included in the final sample. Infants were all born healthy and full-term (after the 37th gestation week or with a birth weight of more than 2.500 g). Twenty-six additional dyads of the original sample (N = 81) were excluded from the neural synchrony analysis due to early termination of the experiment (n = 15), not codable video/audio (n = 5), and bad fNIRS signal quality (n = 6). Subsequent sample sizes were further decreased because parents did either not return the language development questionnaire (n=27) or did not participate in the home visits to assess attachment (n=23). Mothers were, on average, 33.35 years old (SD=4.80 years; range=24-44 years), of European White origin, and highly educated (76% of mothers graduated with a university degree). Participants were recruited from the database of the Department of Developmental Psychology (Wiener Kinderstudien). Most of the registered families were recruited at Vienna General Hospital, and their children were added to the database after their parents gave their written consent. Mothers and infants received a small present for their participation. The local Ethics Committee approved the study (reference no. 00352).

Previous publications, including data from the same sample, comprised analyses on the role of proximity and touch in neural and physiological synchrony [34] and intrapersonal coupling between prefrontal brain activity and respiratory sinus arrhythmia in infants and adults [35].

### Procedure

Mothers and infants participated in three conditions: distal joint watching, proximal joint watching, and free play. In the joint watching conditions, the infant was seated in a highchair (distal joint watching) or in the mother’s lap (proximal joint watching) and watched a calm aquarium video for 90 seconds. The order of distal and proximal joint watching conditions was counterbalanced. In the free play condition, mother and infant were instructed to freely play face-to-face, as they would at home, but without toys and without singing for five minutes. The free play condition always followed the two watching conditions to avoid confounds in the watching conditions. Only interpersonal synchrony data from the free play condition are reported here (see [34] for a comparison of experimental conditions). Mothers’ and infants’ brain activity was measured using functional near-infrared spectroscopy (fNIRS). Their behaviour was micro-coded using ELAN (Version 5.9; Max Planck Institute for Psycholinguistics, The Language Archive, Nijmegen). The dyads’ electrocardiography was also measured but reported on in separate papers [34,35]. Mothers were sent questionnaires on affect, postnatal depression, attachment style, and infant temperament (not reported here) before the first visit to the lab.

At 12 months of age, we visited families at their homes, observed the mother-infant dyad for at least 90 minutes, and rated them on the Attachment Q-Sort [36]. Parents filled out a questionnaire on infants’ expressive language skills (ELFRA-2 [37]) when infants were 24 months old.

### Measures

#### fNIRS

We used two NIRSport 8-8 (NIRx Medizintechnik GmbH, Germany) devices in the tandem setting and simultaneously recorded concentration changes in oxy-haemoglobin (HbO) and deoxy-haemoglobin (HbR) in mother and infant. The 8 × 2 probe sets were attached to an EEG cap with a 10-20 configuration (Figure S1). The probe sets over the left and right inferior frontal gyrus (IFG) surrounded F7 and F8, whereas the probes on the medial prefrontal area (mPFC) surrounded FP1 and FP2. These regions of interest were based on previous work involving adult-child interactions (24,50). In each probe set, eight sources and eight detectors were positioned, which resulted in 22 measurement channels with equal distances of ∼2.3 cm between the infants’ optodes and 3 cm between the mothers’ optodes. The absorption of near-infrared light was measured at the wavelengths 760 and 850 nm, and the sampling frequency was 7.81 Hz.

fNIRS measurements were processed using MATLAB-based functions derived from Homer 2 [38]. Raw data were converted into optical density. Next, optical density data were motion-corrected with a wavelet-based algorithm with an interquartile range of 0.5. Motion-corrected time series were further visually inspected during a quality check procedure (see [39] for further information). Before continuing, we removed 22.87% of the channels from both mother and child from further analyses due to bad signal-to-noise ratio and motion artefacts. Then, slow drifts and physiological noise were removed from the signals using a band-pass second-order Butterworth filter with 0.01 and 0.5 Hz cut-offs. Based on the modified Beer-Lambert Law, we converted the filtered data to changes (μMol) in HbO and HbR. HbR analyses are reported in the supplements.

We assessed the relation between the fNIRS time series in each caregiver and infant using (Morlet) Wavelet transform coherence (WTC) as a function of frequency and time [40]. WTC is more suitable than correlational approaches, as it is invariant to interregional differences in the hemodynamic response function (HRF) [41]. On the other hand, correlations are sensitive to the shape of the HRF, which is assumed to be different between individuals (especially of different ages) and different brain areas. A high correlation may be observed among regions with no blood flow fluctuations. Accordingly, based on previous studies [21], visual inspection, and spectral analyses, the frequency band of 0.063 Hz – 0.167 Hz (corresponding to 6 - 16 s) was identified as the frequency of interest. In the present study, coherence was calculated for each channel combination in 60-second intervals of the interactive free play condition, resulting in 5 (intervals) x 22 (channels) coherence values for each dyad.

Using WTC, we assessed infants’ intra-personal inter-hemispheric functional connectivity between homologous channels in the passive distal watching condition (as an age-appropriate resting phase). The frequency band of interest was determined to be 0.063 Hz - 0.167 Hz (corresponding to 6 - 16 s), according to the potential infant HRF function [42], by visual inspection and spectral analyses. The channels were assigned to the right and left hemispheres, resulting in ten channel combinations connecting both hemispheres. The two channels placed in the middle were excluded for this analysis, as no calculation of inter-hemispheric functional connectivity was possible due to their spatial proximity. Coherence values were calculated per channel combination and then averaged, resulting in one WTC value for each infant.

#### Turn-taking coding

To assess vocal turn-taking in the free play condition (adapted from [1,43]), trained graduate students coded audio recordings of the free-play sessions using ELAN (Version 5.9; Max Planck Institute for Psycholinguistics, The Language Archive, Nijmegen). The experimental sessions were recorded at 25 frames per second. Vocalisations of mother and infant (excluding crying) were micro-coded frame-by-frame. Subsequently, we analysed vocalisations in terms of turn-taking, which is defined as turns that are taken up by others [1]. A turn-taking sequence included two related turns separated by at least 0 seconds or a maximum of 3 seconds. We assessed a) the number of turn-taking sequences from mother to infant (mother vocalisation followed by infant vocalisation), b) the number of turn-taking sequences from infant to mother (infant vocalisation followed by mother vocalisation), c) the number of bi-directional turn-taking sequences (sum of all turn-taking sequences), d) mother response latency, e) infant response latency, f) overall response latencies, g) the number of mother-to-infant overlaps (when infant’s vocalisation overlaps the mother’s preceding vocalisation), h) mother-to-infant overlap duration, i) infant-to-mother overlaps, and j) infant-to-mother overlap duration. Two trained research assistants coded 25% of randomly chosen videos to establish inter-rater reliability. Inter-rater reliability for vocalisations was high (mother vocalisations: K = .98, infant vocalisations: K = .78).

#### Language outcome questionnaire

The ELFRA-2 [37] is a parent-report questionnaire to assess infants’ expressive language skills at two years of age. The questionnaire comprises a list of 260 words, 25 questions on syntax, and 11 questions on morphology. In the present study, we only used the word list to assess infants’ word production at 24 months. Based on the Communicative Development Inventories [44], the list includes 20 categories, such as animals, verbs, and food. The reliability of the items was high, Cronbach’s alpha=0.99.

#### Attachment assessment

We used the Attachment Q-Sort (AQS [36]) to observe the attachment security of the mother-infant dyads at 12 months of age. Two trained research assistants observed the dyad for at least 90 minutes during each home visit and independently evaluated the attachment security. The interrater reliability between raters averaged at r = .75, thus, an averaged AQS score was used in subsequent analyses. These AQS scores ranged between −1.0 and +1.0, with higher scores representing higher attachment security of the observed dyad.

#### Statistical Analysis

All statistical analyses were calculated in RStudio (RStudio Team, 2020). We used Generalised Linear Mixed Models (GLMM), Generalised Linear Models (GLM), and Linear Models (LM) to analyse the hypothesised associations between turn-taking, interpersonal neural synchrony, inter-hemispheric functional connectivity, infant’s language development, and attachment.

Hypothesis 1: Turn-taking and interpersonal neural synchrony over time. Wavelet Transform Coherence (WTC) values were entered as the response variable (assuming a beta distribution and logit link) of a GLMM with interval (1-5), the number of bidirectional turn-taking, response latencies, and ROI (IFG vs lPFC vs mPFC) as fixed factors. We included the interaction between intervals, number of turns, and ROI. ROI was inserted as random slopes and dyads as random intercepts. Random slopes for interval, turn-taking did not lead to model convergence and were thus excluded. To further examine significant main and interaction effects, post-hoc analyses (emmeans) were applied to contrast factors. Multiple comparisons were corrected by using Tukey’s Honest Significant Difference. All continuous predictor variables were z-standardised, and distributions of residuals were visually inspected for each model. Models were estimated using Maximum Likelihood. Model fit was compared using a Chi-Square Test (likelihood ratio test [45]).

Hypothesis 2: Turn-taking and infants’ inter-hemispheric connectivity. We calculated two separate GLM including inter-hemispheric connectivity values as the response variable (assuming a gaussian distribution and logit link). The first GLM tested mother-to-infant response latencies, and number of infant-to-mother turns as the main effects. The second GLM tested the effects of mother-to-infant and infant-to-mother overlaps. Further turn-taking variables are considered in analyses reported in the supplements.

Hypothesis 3: Infants’ expressive language abilities and turn-taking. Infants’ vocabulary at 24 months assessed by the ELFRA questionnaire was entered as the response variable (assuming a gaussian distribution) of three LM. One included the main effects of dyadic turn-taking (number of turns, response latency). Another one included the main effects of infant turn-taking variables (number of turns, response latency, number of overlaps, and overlap duration). The third LM included mother turn-taking variables (number of turns and response latency).

Hypothesis 4: Infants’ attachment and turn-taking. Infants’ attachment security score at 12 months of age assessed by the Attachment Q-Sort was entered as the response variable (fisher’s z-transformed, assuming a gaussian distribution) of two LM. One included the main effects of dyadic turn-taking (number of turns, response latency). Another one included the main effects of infant turn-taking variables (number of turns, response latency, number of overlaps, and overlap duration). The third LM included mother turn-taking variables (number of turns and response latency).

The p-values of the multiple GLMMTMB/GLM/LM were corrected for multiple comparisons when necessary (false discovery rate [46]). We also conducted a power analysis (using a bespoke web app [47]) in relation to the statistical analysis on the sample size of 55. According to previous research [21], we assumed a medium effect size (d=0.5), which yielded 1-ß=0.856. We, thus, deemed the sample size to be appropriate for this set of analyses.

## Results

### Turn-taking patterns

Descriptive statistics for the turn-taking patterns of mother-child dyads (*N*=55) throughout the free play condition are included in Table S1. Number of bidirectional turn-taking and response latencies were highly correlated (turn-taking: β = 0.869, *SE* = 0.068, *t*(53) = 12.78, *p* < .001, response latencies: β = 0.541, *SE* = 0.116, *t*(53) = 4.685, *p* < .001). Neither frequency of turn-taking nor overlaps from mother to infant or infant to mother were significantly related, *p* > .189.

### Turn-taking and interpersonal neural synchrony over time

Firstly, we analysed whether more frequent turn-taking during a proto-conversation was related to mother-infant neural synchrony in frontal regions. The GLMM (*n* = 52) provided a significantly better fit to the data than the null model (χ2 = 28.472, *df* = 11, *p* = .002). The comparison of the full model with the reduced models revealed that the three-way interaction turns*intervals*ROI (χ2 = 11.914, *df* = 2, *p* = .003) and the main effect ROI (χ2 = 11.920, *df* = 2, *p* = .003) were statistically significant. These effects indicate that interpersonal neural synchrony was different between brain regions and that this difference was modulated by the frequency of turn-taking and that their slopes were different over time. Neither the two-way interactions turn-taking*intervals, intervals*ROI, turn-taking*ROI nor the simple effects of turn-taking, intervals and response latencies reached significance (*p* > .139). To understand the three-way interaction, we conducted three separate GLMM to test the interaction between turns and intervals in relation to interpersonal neural synchrony in each ROI. We corrected *p*-values (FDR) for multiple comparisons. The GLMM for the inferior frontal gyrus and lateral prefrontal cortex revealed no significant main effects and interaction effects (*p* > .270). Only the GLMM for the medial prefrontal cortex revealed a significant main effect of turns (*estimate* = 0.005, *SE* = 0.002, 95% *CI* = [0.002 0.010], *z* = 2.742, *p* = 0.006, *p-corrected* = .009), a significant main effect of interval (*estimate* = 0.049, *SE* = 0.019, 95% *CI* = [0.011 0.086], *z* = 2.564, *p* = 0.010, *p-corrected* = .010), and a significant interaction between turns and intervals (*estimate* = -0.002, *SE* = 0.001, 95% *CI* = [-0.003 - 0.0005], *z* = -2.748, *p* = 0.006, *p-corrected* = .009). This significant set of effects shows that more frequent turns were related to higher neural synchrony, specifically in the medial prefrontal cortex.

Neural synchrony between mother and infant also increased over time. But more specifically, more frequent turns were associated with higher neural synchrony in the medial prefrontal cortex in the first intervals (up to the second minute; see *Figure 1*). The relation between turns and neural synchrony is then attenuated over later intervals (third to fifth minute). In addition, we examined whether interpersonal neural synchrony in HbR was to turn-taking but found no significant effects (see Supplements for further details). Next, we explored whether response latencies or overlap frequency were associated with mother-infant neural synchrony. The relations were, however, not significant (*p* > .114).

**Figure 1.**
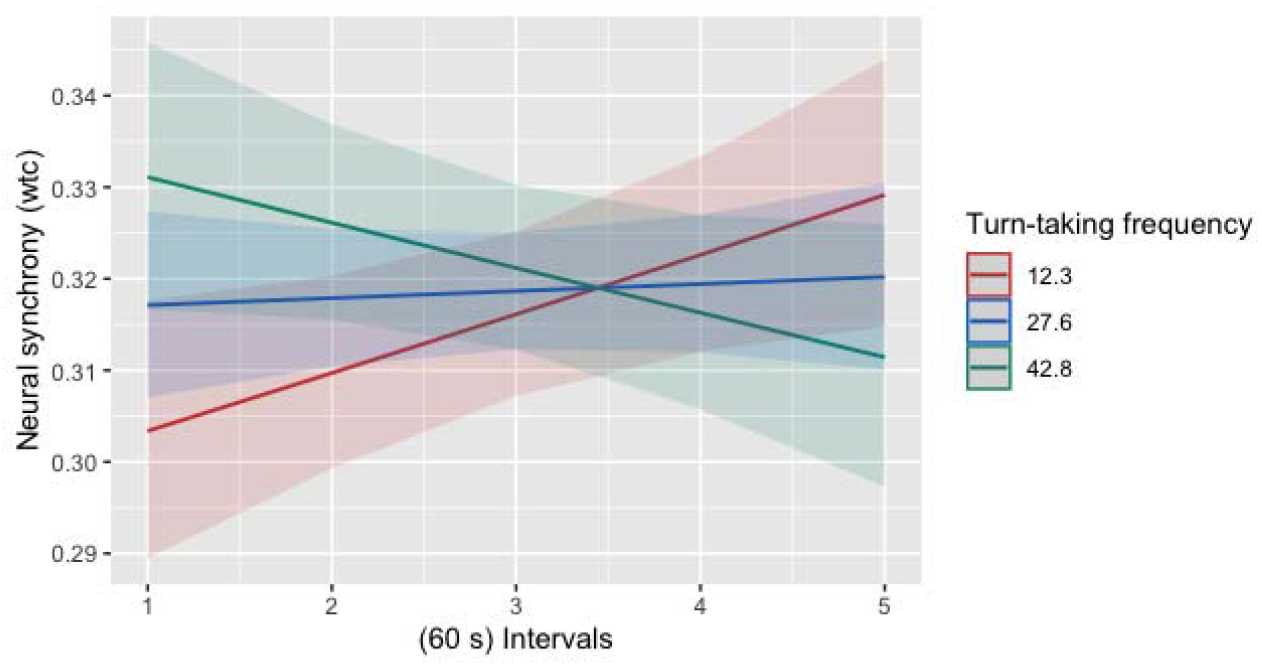
The graph depicts the interaction effect between interval (x Axis) and the number of switching turns (traces, divided in -1 SD=red, estimated mean=blue, +1 SD=green) on interpersonal neural synchrony (measured in wtc; y Axis). A higher turn-taking frequency is associated with higher neural synchrony between mother-infant at the beginning of the proto-conversation. Neural synchrony in later intervals is not associated with turn-taking frequency. The shaded area depicts the 95% confidence interval of the predicted values.

Turn-taking and infants’ inter-hemispheric connectivity. Next, we tested infants’ inter-hemispheric connectivity as a neural correlate to infant-initiated (infant-to-mother) turns and infants’ (mother-to-infant) response latencies. The first model (*n* = 54) output displayed that the frequency of infant-to-mother turns were significantly associated with infants’ inter-hemispheric connectivity (*estimate* = 0.078, *SE* = 0.038, *z* = 2.031, *p* = 0.048), as depicted in Figure 2A. We were able to replicate these results using infants’ inter-hemispheric connectivity in HbR (detailed in Supplements). Mother-to-infant response latencies were however not related to infants’ interhemispheric connectivity (*p* = .701). Moreover, infants’ and mother’s frequency of overlaps were not significantly associated with inter-hemispheric connectivity (*p* > .255). Taken together, more infant-initiated turns in the proto-conversation were associated with higher inter-hemispheric connectivity in infants’ brains.

**Figure 2.**
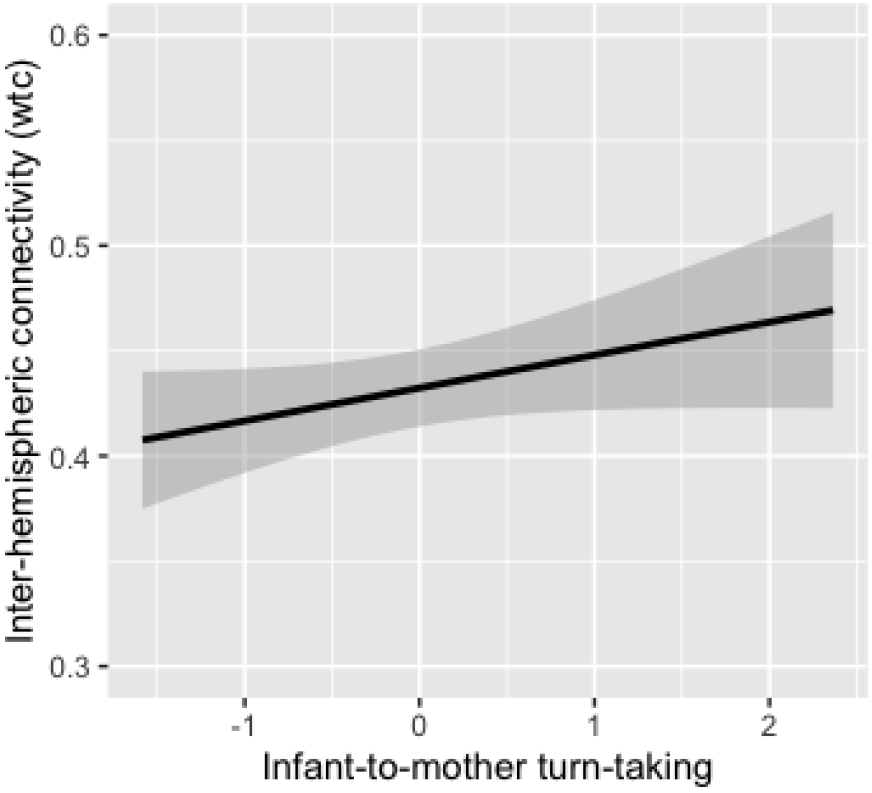
Infant-to-mother turn-taking (x Axis) is positively associated with infants’ inter-hemispheric connectivity (assessed using wtc; y Axis). The predicted values are plotted in black, and the shaded area depicts the 95% confidence interval.

Infants’ expressive language abilities and turn-taking. We tested the relation between infants’ vocabulary at 24 months and dyadic as well as individual turn-taking patterns during the mother-infant interaction at 4-6 months (*n* = 28). The first LM assessed the relation between infants’ vocabulary and dyadic turn-taking patterns and revealed a significant correlation between the number of turns of the dyad and infant vocabulary (*estimate* = 0.137, *SE* = 0.021, *z* = 6.509, *p* < .001). More frequent turn-taking were related to infants’ larger vocabulary at 24 months (*Figure 3A*). Next, we split the dyadic turns into uni-directional turns and ran an additional regression analysis. The result showed that the number of mother-to-infant turn-taking explained more variance in infants’ vocabulary than infant-to-mother turn-taking (*estimate* = 0.164, *SE* = 0.030, *z* = 5.414, *p* < .001). The third LM assessed the relation between infants’ vocabulary and mother-to-infant overlap frequency and found a significant negative relation (*estimate* = -0.066, *SE* = 0.021, *z* = -3.151, *p* < .001). Fewer mother-to-infant overlaps were associated with infants’ larger vocabulary (*Figure 3B*). Infants’ vocabulary was not significantly related to infant-to-mother overlaps (*p* = .794).

**Figure 3.**
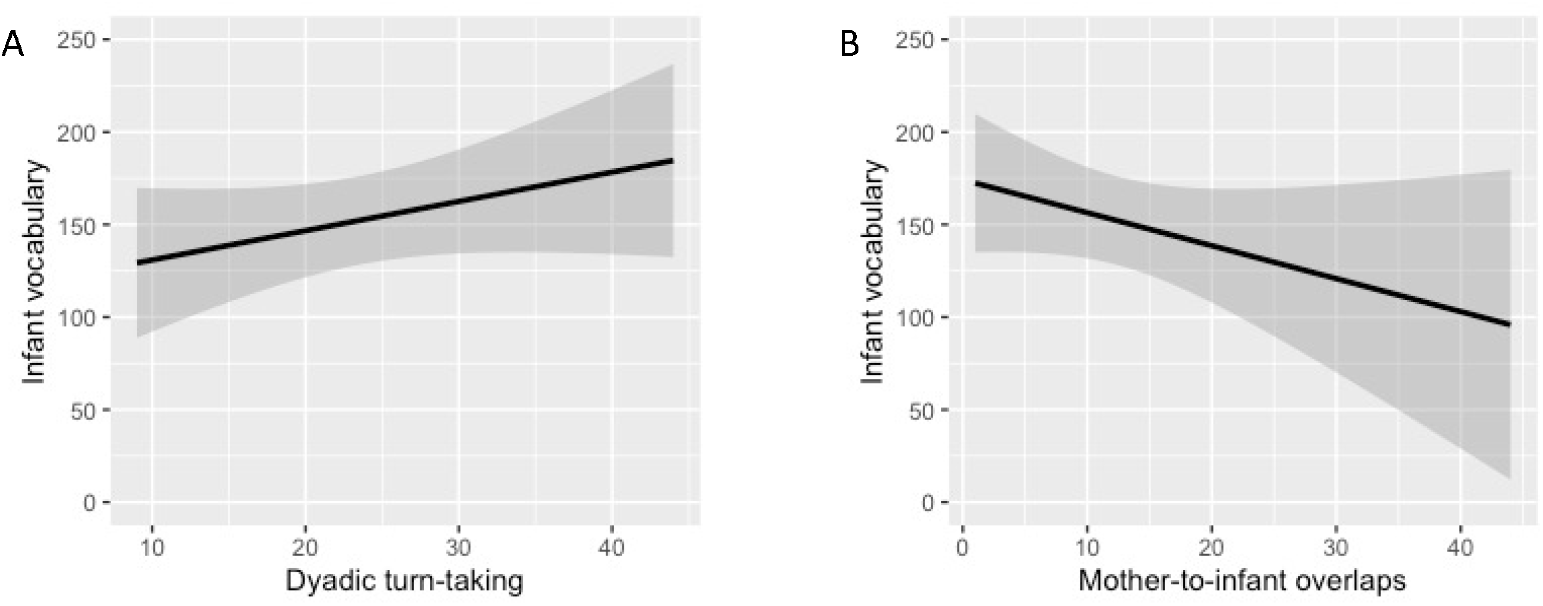
(A) The graph depicts the relation between dyadic turn-taking frequency (x Axis) at 4-6 months of age and infants’ vocabulary at 24 months of age (y Axis). (B) The graph depicts the relation between infant overlaps (x Axis) at 4-6 months and infants’ vocabulary at 24 months of age (y Axis). Predicted values are plotted in black, and the shaded area depicts the 95% confidence interval.

Infants’ attachment and turn-taking. Next, we tested the relation between mother-infant turn-taking patterns at 4-6 months and infants’ attachment security at 12 months. Attachment security was not related to mother-infant turn-taking patterns, *p-corrected* > .122. In an exploratory analysis, we dichotomised infants’ attachment scores into securely and insecurely attached infants (split at *r* = 0.3 as per [36]). The tested relation between secure vs insecure attachment and turn-taking patterns was not significant, *p-corrected* > .140.

## Discussion

In the present study, we examined turn-taking patterns in mother-infant proto-conversations in relation to interpersonal neural synchrony and developmental outcome measures: infants’ inter-hemispheric connectivity, expressive language abilities, and attachment. The more turn-taking mother and infant showed the more they displayed interpersonal neural synchrony during the proto-conversation. The relation was stronger in the initial phase of the interaction and decreased over time. Both infants’ number of turns and overlaps (in response to the mothers’ vocalisation) were related to higher inter-hemispheric connectivity in infants’ frontal brain regions. The number of turns in general, but especially infants’ turns, was related to a larger vocabulary at 24 months of age.

However, mother-infant turn-taking was not related to the infant’s attachment security. The results underscore the role of interpersonal neural synchrony in early turn-taking as well as brain maturation and language development as potential outcomes related to coordinated proto-conversation in early caregiver-infant interactions.

As predicted, we observed a positive relation between the frequency of turns and neural synchronisation between mothers and their infants in a proto-conversation. This association was limited to the medial prefrontal cortex among our regions of interest. Medial prefrontal regions have been implicated in caregiver-infant synchrony, especially in face-to-face exchanges [48]. This is consistent with the role of the medial prefrontal cortex in processing communicative signals directed to the self and, more generally, in social cognition and mentalising processes [49–51]. Synchronisation of rhythmic brain activities in this region could be linked to high levels of mutual engagement, thus facilitating mutual prediction and fluid turn-taking between interaction partners. Also, in line with our predictions, neural synchrony in the mother-infant dyads generally increased over the course of the free play exchange. However, the positive association between frequency of turns and neural synchrony was unexpectedly limited to the early phases of the mother-infant interaction.

Our hypothesis that neural synchrony might be a shared underlying mechanism for turn-taking across earlier and later human development can therefore neither be fully confirmed nor dismissed. In contrast to the current results, neural synchrony increased over time in mothers and their preschool-aged children in association with more frequent turn-taking in a previous study [21]. There are several possible interpretations for the partial discrepancy between the current findings and these earlier results with older children. While neural synchrony might help mother-infant dyads getting attuned to each other early on during a proto-conversation, upholding a preverbal exchange with frequent turns might not rely on persistent neural synchrony in the same way as might be the case in later (more complex) verbal conversations. Alternatively, the oscillator model of turn-taking [13] might indeed be a universal mechanism for precisely timed turn-taking, but we were not able to capture this over the course of the proto-conversations here, e.g., due to our focus on slower hemodynamic brain responses instead of temporally better resolved electroencephalography. Future research could elucidate the functional link between neural synchrony and turn-taking in different age groups further by experimentally manipulating turn-taking and measuring resulting effects on neural synchrony and vice versa, by manipulating neural synchrony through neurofeedback or transcranial stimulation methods [52].

Next, we examined the relationship between mother-infant turn-taking and infant maturity at the neural level, measured by inter-hemispheric connectivity. Consistent with our hypothesis, we found that more frequent infant-to-mother turn-taking, but not mother-to-infant response latencies, was positively related to brain maturation. These results support previous findings that infants’ individual differences in inter-hemispheric connectivity can be identified and related to individual emotional and sensorimotor differences [27,28]. Interestingly, at 4-6 months, more mature infants might more likely engage in early proto-conversations and express themselves vocally, independent of how well coordinated their own vocalisations might be [30,53]. In sum, our results show that the frequency of proto-conversational patterns (i.e., turn-taking), independent of their quality and thus response latency, is connected to infant brain maturation. The direction of this link is, however, unclear as we assessed turn-taking and inter-hemispheric connectivity at the same time point.

Even though early social interactions are maintained to be a core component of children’s language development [10], few studies have explicitly evidenced a relation between early turn-taking and infants’ language abilities. Here, we find that more frequent turn-taking between mothers and infants as young as 4-6 months of age, especially mother-to-infant turns, is related to their expressive vocabulary skills at 24 months. Our results extend recent findings showing that turn-taking in children between 2 and 24 months of age and their parents is related to their vocabulary growth [31] and emerging communicative capacity [29]. Not only do the number of overall turns matter, but the results again point to the integral and contingent response of the caregiver as a factor relating to infants’ language skills. Importantly, frequent (mother-to-)infant overlaps during the proto-conversation were negatively related to infants’ later vocabulary. We suspect that the negative relation between overlaps and infants’ language development might reflect infants’ drastic reduction of overlap frequency and duration during proto-conversations as they grow [1]. Even though call overlapping is discussed as an important communicative signal in several species [54], mostly, overlap avoidance is discussed in relation to optimized human communication (e.g.,[7]). Vocal matching and vocal imitation in human-mother infant interactions occurs less often [55] and seem to be more relevant for mother-infant musical interactions [56]. Overlap avoidance becomes even more important when infants start to speak their first words, implicating infants’ inhibitory and predictive abilities to coordinate with others as relevant for infants’ language development [10]. Consistent with previous results, response latencies were again not significantly related to infants’ expressive vocabulary at 24 months. This finding indicates that infants accept a range of latencies, especially by their caregivers, when engaging in a proto-conversation [57]. Taken together, the results highlight the importance of emerging turn-taking in frequency but not latency in caregiver-infant proto-conversations when preparing preverbal infants for their first words. While we focussed exclusively on vocabulary size, future studies might consider including other potentially relevant variables, such as syntax development or pragmatics, to gain a more comprehensive view on the relation between turn-taking and language development.

Next to supporting the development of infants’ language skills, conversational turn-taking is suggested to help build connections between people [11,22]. Previous findings indicate that specifically insecurely attached children showed longer response latencies during caregiver-child proto-conversations [32]. However, we did not detect a significant relation between early preverbal turn-taking patterns and infant attachment security. The lack of a significant association could be due to several reasons: Firstly, we assessed very early forms of turn-taking and attachment at a much younger age, while the previous study assessed older infants and toddlers. The relation between turn-taking and attachment could thus develop as infants grow beyond the preverbal phase, with infants’ attachment representations maturing [58]. Secondly, our sample might have been too homogenous for infants’ turn-taking abilities to differ significantly between groups of securely and insecurely attached children. Overall, our results indicate that future studies might need to include a larger sample of insecurely attached infants to precisely examine early turn-taking as a way to facilitate the connection between caregiver and child.

Our results highlight the importance of preverbal turn-taking between caregivers and infants in early development. Turn-taking arises together with interpersonal neural synchronisation in the medial prefrontal cortex, a potential biomarker of mutual engagement and prediction. Moreover, turn-taking in proto-conversations is related to infants’ brain maturation and language development. Our results also indicate that the range of turn-taking qualities in the present sample might comprise “good enough” communication patterns to facilitate the bond between healthy and (mostly) securely attached infants and their caregivers [12]. Therefore, a future research avenue could include investigating samples that show altered interaction patterns, such as insecurely attached infants or anxious caregivers [7,59].

Taken together, our results show that mother-infant turn-taking, similar to later in development [21], relies on a high level of mutual engagement. More mature infants initiate more turns and more responsive mothers have infants with better language outcomes. In addition, bidirectional turn-taking frequency is related to higher levels of neural synchrony, pointing to the intricate link between both partners being mutually engaged with one another (to establish neural synchrony) and a fluid turn-taking exchange. Yet, the directionality of that link and thus mechanistic underpinnings to turn-taking remain to be tested.

## Acknowledgements

This research was funded by the University of Vienna and by a doctoral stipend from the Studienstiftung des Deutschen Volkes granted to T.N. We are grateful to Liesbeth Forsthuber, our student assistants Anja Lueger, Melanie Huber, and Hannah Meller for their support in data acquisition, video, and attachment coding. Additionally, we thank all families who participated in the study and the Department of Obstetrics and Gynaecology of the Vienna General Hospital for supporting our participant recruitment.

