## Supplementary material for "Your turn, my turn. Neural synchrony in mother-infant proto-conversation": Table S1

**Turn-taking over time.** We also examined whether turn-taking patterns changed over time. We, therefore, divided the free play interaction into five x one-minute epochs and analyzed the change in frequency of turns and response latencies using linear mixed-effects models. However, both models revealed no significant changes in neither turn-taking nor response latencies over time, *p* > .350.

**Turn-taking and interpersonal neural synchrony (HbR) over time.** Here, we analysed whether more frequent turn-taking during a proto-conversation was related to mother-infant neural synchrony of HbR in frontal regions. We tested the same GLMM as for interpersonal neural synchrony as in HbO. The GLMM (n=52) showed no better fit than the null model, *p* = .283 and none of the tested effects (time, region and turn-taking) were significant, *p* > .123.

**Turn-taking and infants’ inter-hemispheric connectivity (HbR).** Next, we tested infants’ inter-hemispheric connectivity in HbR as a neural correlate to infants’ turn-taking patterns. The HbR Model confirms the finding from HbO Models. The frequency of mother-to-infant turns were significantly associated with infants’ inter-hemispheric connectivity in HbR (*estimate* = 4.729, *SE* = 0.953, *z* = 4.962, *p* < 0.001, *p-corrected* < .001). However, we did not find further significant relations between neither infant-to-mother turns or infant overlaps and inter-hemispheric connectivity, *p* > .212.

**Table S1.** Descriptive statistics on turn-taking

| Variables | M | SD | min | max |
| --- | --- | --- | --- | --- |
| Turns | 27.85 | 14.58 | 3 | 62 |
| Mother-to-infant turns | 13.82 | 7.48 | 1 | 30 |
| Infant-to-mother turns | 14.04 | 7.60 | 2 | 32 |
| Response latencies (ms) | 644.7 | 233.42 | 217.00 | 1338.79 |
| Mother-to-infant response latencies (ms) | 649.48 | 281.12 | 120.00 | 1266.67 |
| Infant-to-mother response latencies (ms) | 639.91 | 249.32 | 224.00 | 1336.00 |
| Mother-to-infant overlaps | 12.58 | 8.21 | 1 | 44 |
| Mother-to-infant overlap durations (ms) | 1197.91 | 578.56 | 300 | 3426.67 |
| Infant-to-mother overlaps | 8.05 | 5.95 | 0 | 22 |
| Infant-to-mother overlap durations (ms) | 1352.97 | 720.30 | 0 | 3666.67 |

**Figure S1.** (A) The graph depicts the Wavelet Transform Coherence calculations between homologous fNIRS channels in mother (left) and infant (right) dyads. (B) The graph shows the intrapersonal channels and connections used to calculate inter-hemispheric connectivity in infants.

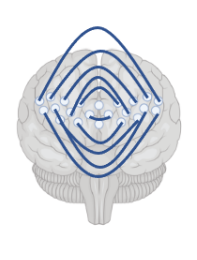

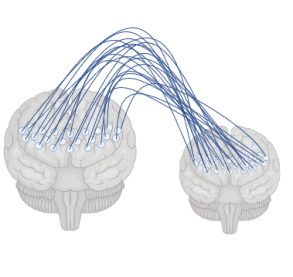

A

B

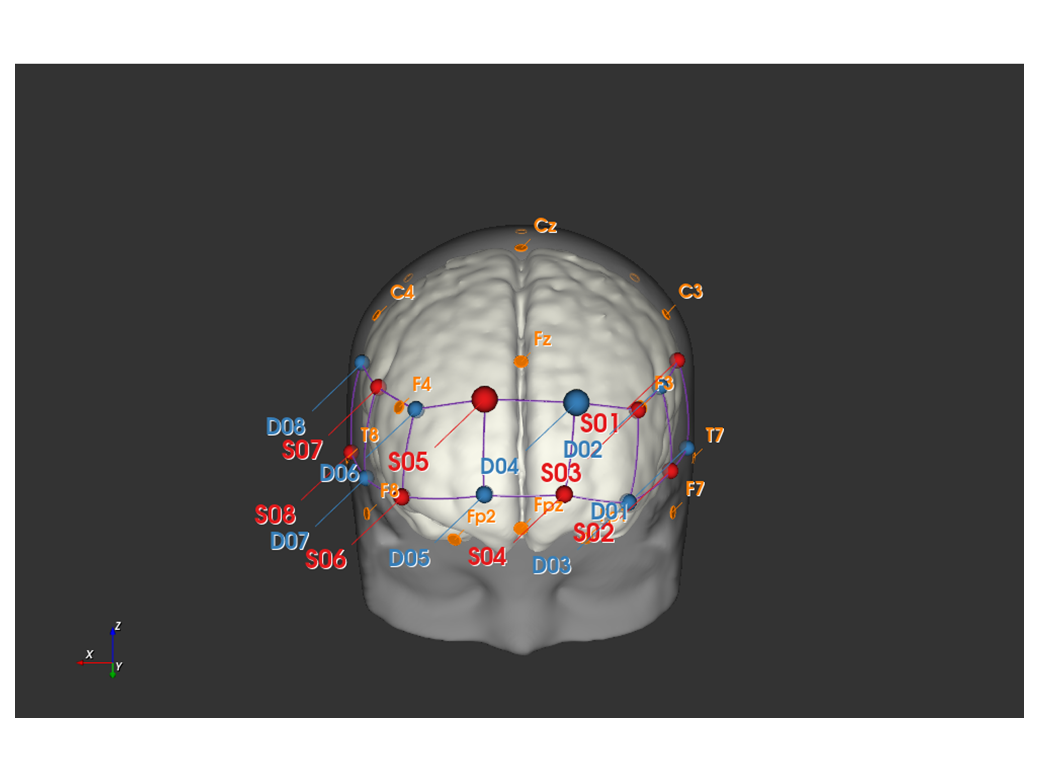

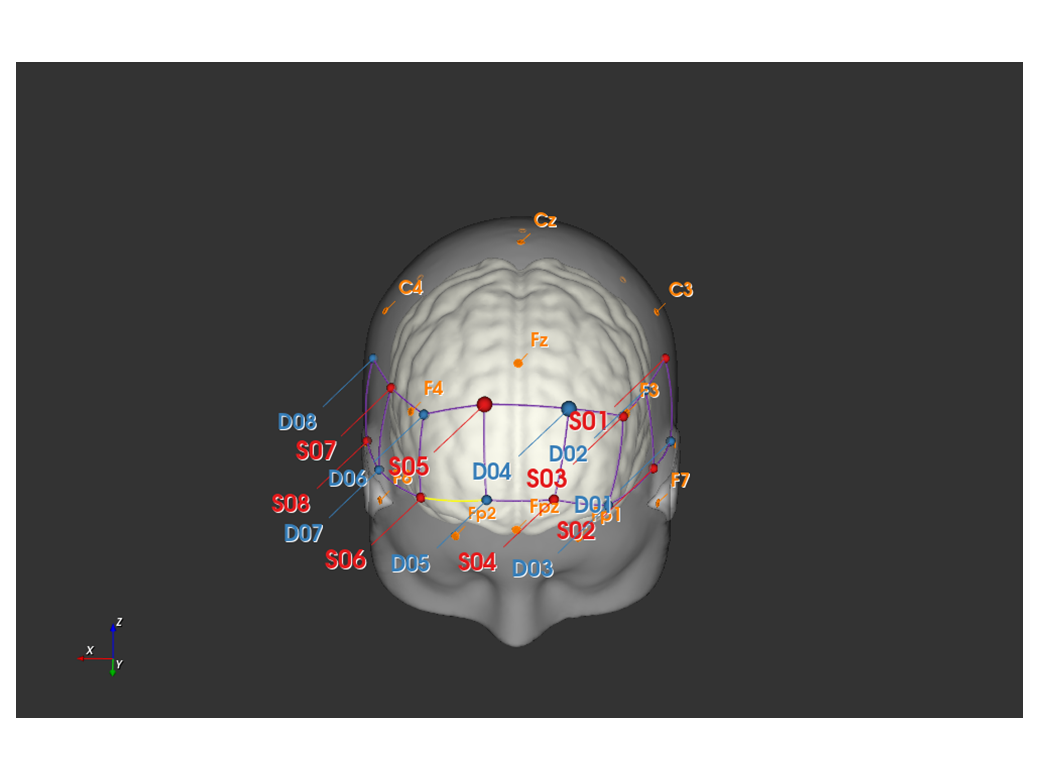

**Figure S2.** Optode positions are projected onto a 2-5-month-old infant head model (Infant Atlases 0-4.5 years; top) and an adult head model (ICBM 152 Nonlinear atlases version 2009; bottom). EEG 10/20 reference labels, which we used to place optode positions onto the caps, are marked in orange. Sources are in red, detectors in blue. Projections were implemented using NIRSite 2021.4.
